# Chromatin Accessibility Maps Provide Evidence of Multilineage Gene Priming in Hematopoietic Stem Cells

**DOI:** 10.1101/2020.11.24.394882

**Authors:** Eric W. Martin, Jana Krietsch, Roman E. Reggiardo, Rebekah Sousae, Daniel H. Kim, E. Camilla Forsberg

## Abstract

Hematopoietic stem cells (HSCs) have the capacity to differentiate into vastly different types of mature blood cells. The epigenetic mechanisms regulating the multilineage ability, or multipotency of HSCs are not well understood. To test the hypothesis that *cis* regulatory elements that control fate decisions for all lineages are primed in HSCs, we used ATAC-seq to compare chromatin accessibility of HSCs with five unipotent cell types. We observed the highest similarity in accessibility profiles between Megakaryocyte Progenitors and HSCs, whereas B cells had the greatest number of regions with *de novo* gain in accessibility during differentiation. Despite these differences, we identified *cis* regulatory elements from all lineages that displayed epigenetic priming in HSCs. These findings provide new insights into the regulation of stem cell multipotency, as well as a resource to identify functional drivers of lineage fate.

**HIGHLIGHTS:**

- HSCs have higher global chromatin accessibility than any unilineage progeny
- Megakaryocyte Progenitors are the most closely related unipotent cell type to HSCs
- B cell commitment involves *de novo* chromatin accessibility
- Evidence of *cis* element priming of lineage-specific genes in HSCs

## INTRODUCTION

Multipotency is a key feature of hematopoietic stem cells (HSCs) and essential for their ability to produce all types of blood and immune cells *in situ* and upon therapeutic stem cell transplantation. The mechanistic basis of multipotency is unclear, but previous studies have shown that the regulation of differentiation programs is achieved, in large part, through epigenetic remodeling of *cis*-regulatory elements (CREs) (Forsberg et al., 2000; Shivdasani et al., 1997; Wang et al., 2015). Thus, HSC multipotency may be enabled by accessible non-promoter CREs that keep loci competent for transcription factor binding and gene activation without active expression. Such selective “CRE priming” may underlie the developmental competence of specific cell types, which is then acted upon by inductive signals to gradually specify fate (Waddington, 1940). When all CREs that drive differentiation and lineage choice are primed in stem cells, that stem cell is in a permissive state **(Figure 1A**) and is competent to initiate differentiation into all mature lineages.

**Figure 1:**
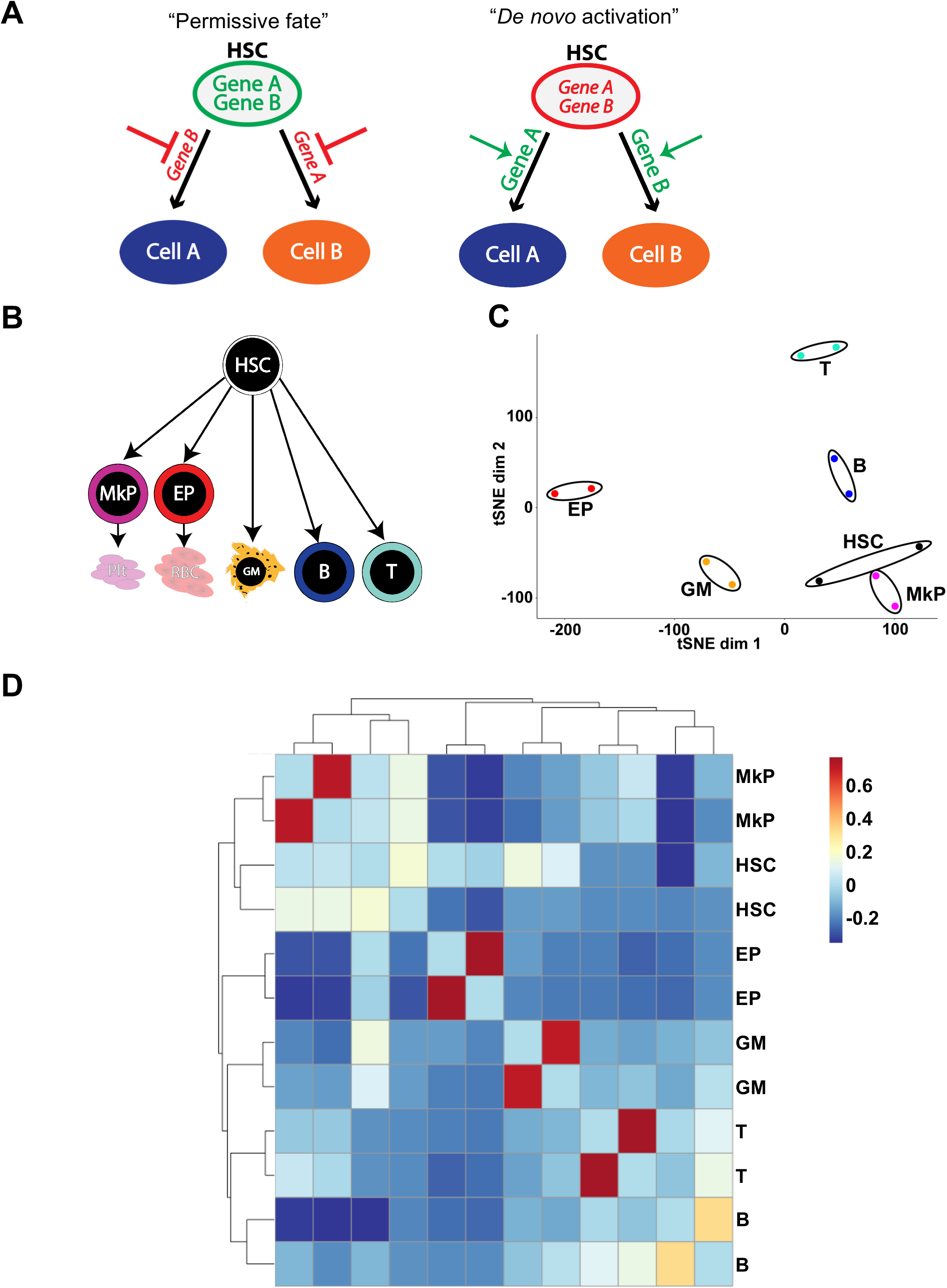
ATAC-seq maps of hematopoietic cell populations exhibit a high degree of reproducibility between replicates and a tight association of MkPs to HSCs. **A)** Two models of epigenetic regulation of HSC fate. In the “permissive fate” model, CREs of lineage-specific genes of all possible lineage outcomes are in an accessible state (green) in HSCs, keeping genes “primed” for subsequent activation. After lineage commitment occurs towards one fate, the accessibility of primed elements of the alternative fate is restricted by epigenetic remodeling (red). In contrast, the “*de novo* activation” model posits that CREs of lineage-specific genes are in an inaccessible state (red) in HSCs, keeping genes silenced in heterochromatin. Lineage commitment occurs by *de novo* decondensing of chromatin at the appropriate CRE, allowing for subsequent activation of the differentiation program (green). The CREs of alternative lineage fates remain restricted as heterochromatin (red). **B)** Schematic diagram of the hematopoietic cells used in this study. Six cell populations were investigated: multipotent HSCs (Hematopoietic Stem Cells), unilineage MkPs (Megakaryocyte Progenitors) and EPs (Erythroid Progenitors), and mature GMs (Granulocyte/Macrophages), B cells, and T cells. **C)** tSNE analysis of the ATAC-seq peaks revealed a high concordance of biological replicates. MkPs clustered close to HSCs, while EPs, GMs, B, and T cells separated across the tSNE plot. **D)** Hierarchical clustering revealed high concordance of cell type-specific replicates. Similar to the tSNE analysis, MkPs clustered closest to HSCs. B and T cells were closely associated to each other but distant to HSCs, while GMs and EPs were contained within their own branches, closer to HSCs.

We sought to test two models of HSC multipotency that are based on regulation of chromatin organization: the “permissive fate model” and a “*de novo* activation model” **(Figure 1A)**. Supporting a role for the permissive model in stem cell lineage potential are observations of bivalent histone domains that maintain key developmental genes in embryonic stem cells (ESCs) poised for activation (Bernstein et al., 2006), and an overall accessible chromatin state in both ESCs and HSCs compared to lineage-restricted progenitors and mature cells (Gaspar-Maia et al., 2009, 2011; Ugarte et al., 2015). When differentiation occurs, the genes poised for differentiation into the induced lineage are activated while CREs that would drive differentiation into alternative lineages are silenced into heterochromatin. This has been observed in ESCs and during differentiation of ESCs into endoderm (Wang et al., 2015; Xu et al., 2009). Our observation of global chromatin condensation and localization of H3K9me3-marked heterochromatin towards the nuclear periphery during HSC differentiation also support the permissive model (Ugarte et al., 2015). Inversely, in the *de novo* activation model **(Figure 1A)**, CREs that drive lineage fate are inaccessible in HSCs. Differentiation and lineage choice occur by “unlocking” these CREs. Transcriptional and functional analyses of hematopoietic stem and progenitor cells support this *de novo* model, where lymphoid potential is *gained* in progenitor cells rather than being a consequence of CRE priming in HSCs (Boyer et al., 2019; Cabezas-Wallscheid et al., 2014; Forsberg et al., 2005; Månsson et al., 2007).

In order to interrogate these models and how they pertain to the regulation of competence in hematopoiesis, as well as gain a better understanding of the relationships between epigenetic, transcriptomic and functional observations, we mapped global chromatin accessibility using the Assay for Transposase Accessible Chromatin by High Throughput Sequencing (ATAC-seq) (Buenrostro et al., 2013). This assay allows assessment of high resolution, genome-wide chromatin accessibility throughout differentiation programs of rare cells. The dynamics of chromatin accessibility in erythro-megakaryopoiesis (Heuston et al., 2018) and granulocyte/macrophage development (Buenrostro et al., 2018) have been highly informative. From these studies, the bulk observations gave us insight into the dynamics of lineage commitment during hematopoiesis, while single-cell analysis revealed the heterogeneity of epigenomic states and, therefore, lineage bias in progenitors throughout hematopoiesis. Based on those studies, as well as reports of global chromatin accessibility of embryonic (Bernstein et al., 2006; Bulut-Karslioglu et al., 2018; Gaspar-Maia et al., 2011) and hematopoietic (Cabal-Hierro et al., 2020; Ugarte et al., 2015) stem cells, we hypothesized that HSCs are in a permissive chromatin state where CREs that control fate decisions are primed in HSCs. Here, we tested this hypothesis by performing in-depth ATAC-seq investigation of HSCs and 5 unipotent lineage cell populations representing the five main hematopoietic lineages **(Figure 1B)**, as defined by previously published phenotypes (Boyer et al., 2011, 2019).

## RESULTS

### Mapping of chromatin accessibility in HSCs and unipotent lineage cells identified a tight association of MkPs to HSCs

To determine the dynamics of genome accessibility throughout hematopoiesis, we sorted six primary hematopoietic cell types (**Figure 1B**) and performed ATAC-seq. We identified 70,731 peaks in HSCs, 47,363 peaks in MkPs, 38,007 in EPs, 30,529 in GMs, 70,358 in B cells, and 51,832 in T cells **(Table 1)**. From these peak-lists we combined and filtered the peaks to only the most significant peaks using the chromVAR package (Schep et al., 2017) and identified a total of 84,243 peaks, referred to as the master peak-list throughout the study **(Table 1)**. To assess data quality, we analyzed replicate clustering and cell type relationships of all 6 cell types using principal component analysis and dimensionality reduction as a t-Distributed Stochastic Neighbor Embedding (tSNE) plot (Schep et al., 2017). All biological replicate samples closely associated with each other by tSNE analysis (**Figure 1C**), as well as by hierarchical clustering using the chromVAR output (**Figure 1D**). We observed two primary clusters in **Figure 1D**: an HSC/MkP cluster and all other cell types. We also observed a distinct lymphoid cell subcluster containing only B and T cells, while GMs and EPs clustered independently. MkPs have the most similar accessibility to HSCs, with the ranking of the other cell types from most to least similar as EPs, GMs, Bs and then Ts. This is consistent with our tSNE analysis (**Figure 1C**), where HSCs and MkPs closely associated with each other, and with studies that have reported a close relationship of HSCs with the megakaryocyte lineage (Carrelha et al., 2018; Rodriguez-Fraticelli et al., 2018) and that erythropoiesis requires chromatin remodeling for differentiation to occur (Heuston et al., 2018).

**Table 1:** Peak counts and peak distribution relative to protein-coding gene promoters in each cell type.

| Cell Type | ATAC peaks | Promoter Peaks ($\pm 500$bp of TSS) | non-promoter peaks | | |
| --- | --- | --- | --- | --- | --- |
|  |  |  | coding (exons+TTS+TSS) | Introns | Intergenic |
| Master Peak-list | 84,243 | 13,171 | 5,243 | 34,137 | 31,692 |
| HSC | 70,731 | 27,973 | 4,166 | 18,931 | 19,661 |
| MkP | 47,363 | 23,998 | 2,013 | 10,036 | 11,316 |
| EP | 38,007 | 23,243 | 2,014 | 7,040 | 5,710 |
| GM | 30,529 | 15,559 | 1,440 | 6,697 | 6,833 |
| B | 70,358 | 24,596 | 4,461 | 21,210 | 20,091 |
| T | 51,832 | 25,103 | 2,016 | 11,929 | 12,784 |

### Visualization and comparison of ATAC-seq data generated in this study correlated with known expression patterns of cell type-specific genes

As another assessment of the quality and reproducibility of our ATAC-seq data, we visualized the ATAC-seq signals across promoters of genes with known cell type-specific expression patterns, plus a negative (expressed in none of the cell types) and a positive (expressed in all of the cell types) control, using the Gene Expression Commons (GEXC) expression database (Seita et al., 2012): *Gapdh* (expressed in all cell types)*, Fezf2* (not expressed in any cell type)*, Klf1* (EPs only)*, Gp6* (MkPs only)*, Ly6g* (GMs only), *CD19* (B cells only), and *Ccr4 (*T cells only) (**Figure 2A,B**). *Ly6g* was not available in GEXC but is a well-known GM-selective gene (Hestdal et al., 1991). We observed the expected accessibility peaks in each cell type, as well as a minimal signal from cell types without expression of those genes (**Figure 2B**). The overall high level of reproducibility between independent sample replicates and clustering strategies (**Figure 1C,D**), as well as the expected accessibility in cell type-specific genes (**Figure 2**), indicated that we had generated high-quality chromatin accessibility maps of these 6 cell types.

**Figure 2:**
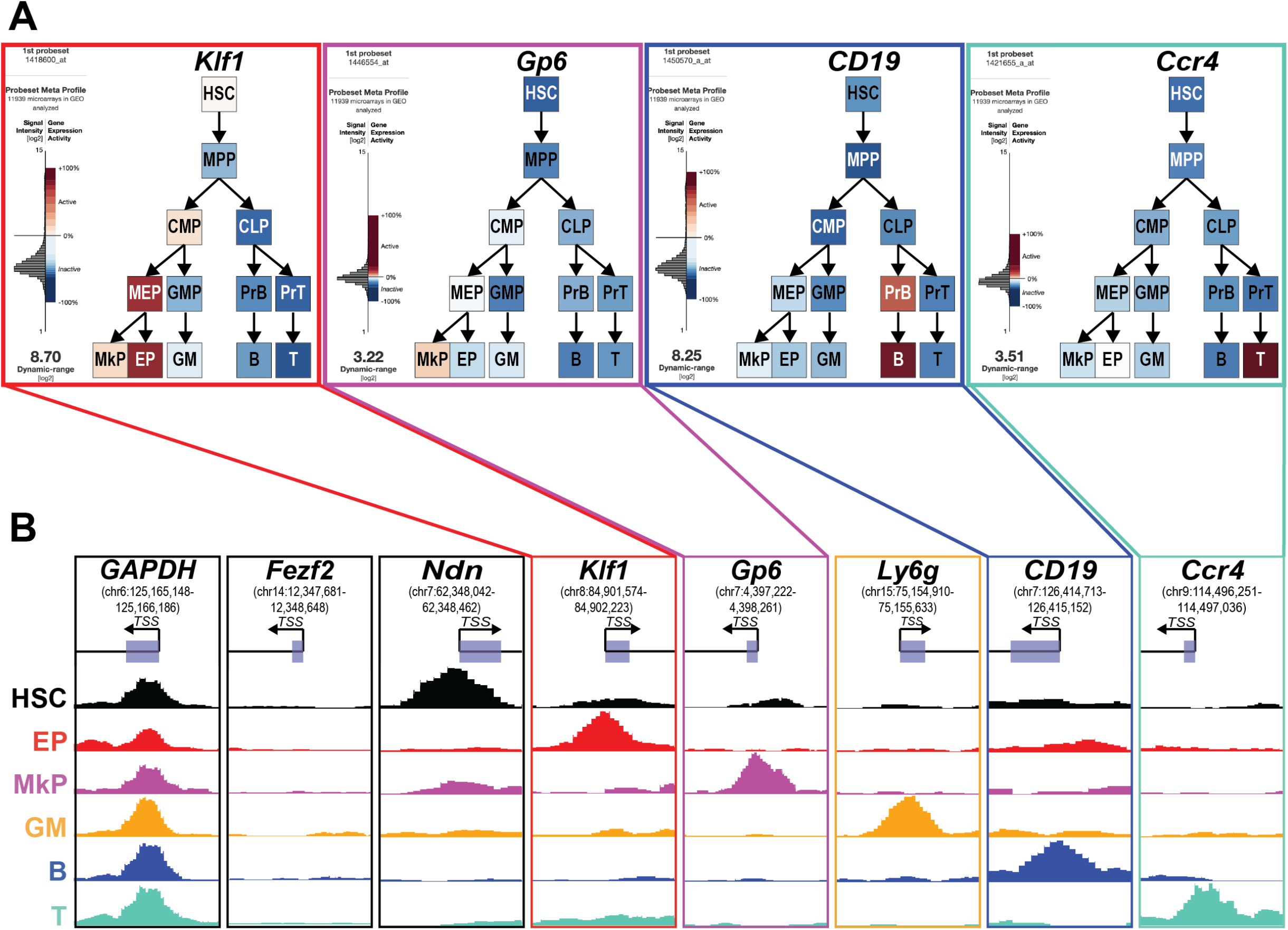
Promoter accessibility correlated with known expression patterns of cell type-specific genes. **A)** Lineage-specific expression of one example gene each for MkPs, EPs, B, or T cells. The level of expression (red=high; blue=low/not expressed) according to the Gene Expression Commons (GEXC) database of an example gene with cell type-specific ATAC-seq promoter peak. The probeset for the GM-specific *Ly6g* is not present in GEXC and therefore not displayed. **B)** Cell type-specific chromatin accessibility visualized as ATAC-seq read counts at transcription start sites (TSS) using UCSC Genome Browser snapshots. Depiction of the six ATAC-seq libraries used in this study with example genes that had ATAC-seq signal in all samples (*GAPDH*; positive control), no samples (*Fezf2*; negative control), or in a specific cell type: HSCs (*Ndn*), EPs (*Klf1*), MkPs (*Gp6*), GMs (*Ly6g*), B cells (*CD19*), and T cells (*Ccr4*).

### HSCs have greater global accessibility and undergo more extensive chromatin remodeling upon lymphoid differentiation

Using a number of quantitative, but non-sequence-specific assays, we previously reported that chromatin is progressively condensed upon HSC differentiation into unilineage and mature cells (Ugarte et al., 2015). To test whether the ATAC-seq data recapitulated these findings, we quantified the total number of distinct peaks, as well as the cumulative read-counts for all peaks, for each cell type. First, we took each cell type’s optimal peak-list from the Irreproducible Discovery Rate (IDR) analysis (Li et al., 2011) and reported the number of peaks. We observed the highest number of peaks in HSCs (**Figure 3A**), closely followed by B cells. In parallel, we quantified global accessibility by calculating the normalized average signal over the master peak-list for each cell type by generating histograms using HOMER (Heinz et al., 2010). We observed similar ordering compared to the peak number, with HSCs having the highest average signal and B cells the second highest (**Figure 3B**). The low signal in EPs is possibly due to widespread transcriptional silencing as the next step towards becoming highly specialized red blood cells and ejection of nuclei (An et al., 2014). Although these measurements are not completely independent, HSCs displayed both the highest number of peaks and the greatest peak signal. These results are consistent with our previous findings of progressive chromatin condensation upon HSCs differentiation (Ugarte et al., 2015).

**Figure 3:**
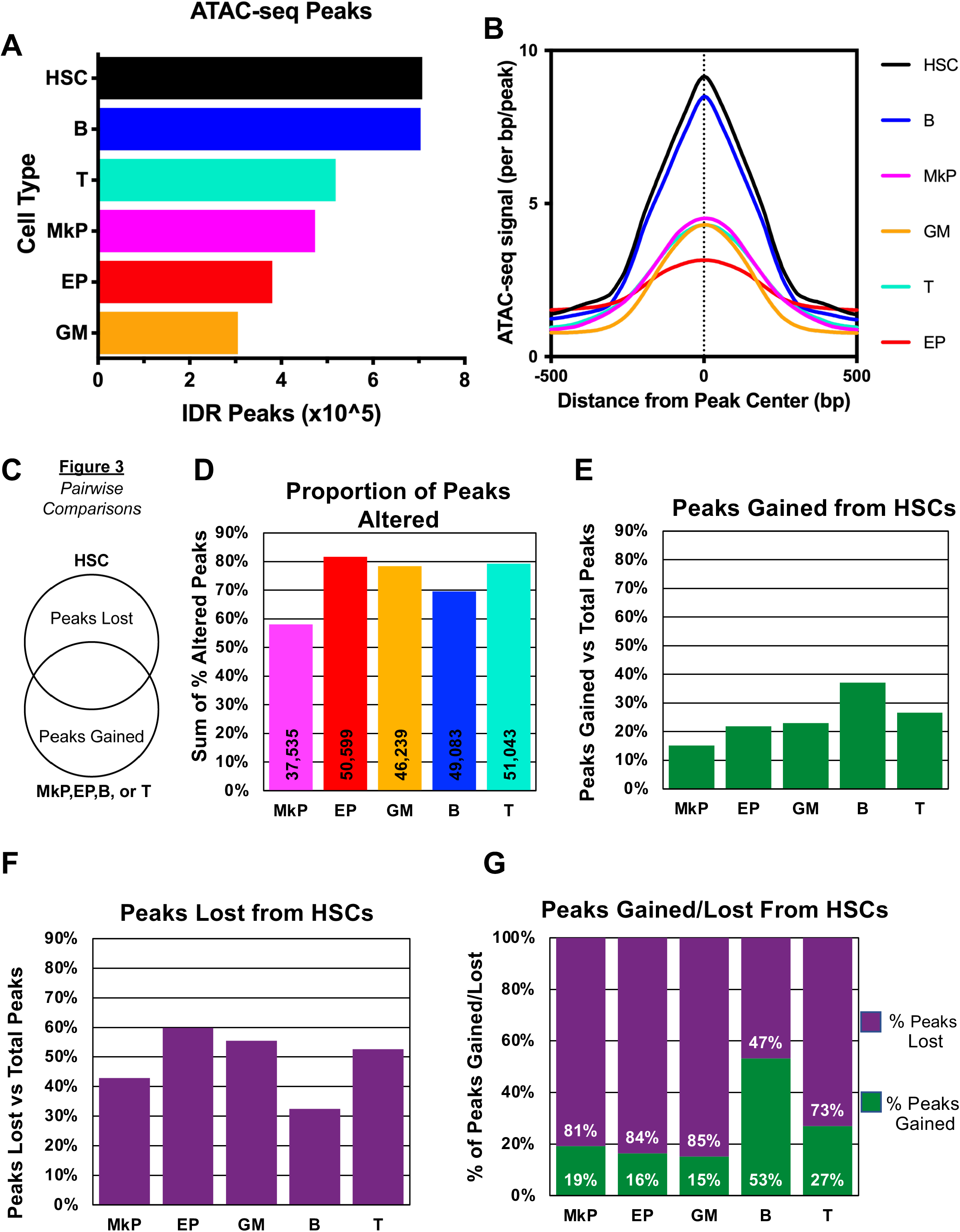
Greater overall global accessibility of HSCs and more extensive chromatin remodeling upon lymphoid differentiation. **A)** HSCs had the highest number of peaks of all hematopoietic cell types. The total number of peaks are displayed. HSCs had the highest number of peaks followed by B cells, T cells, MkPs, EPs, then GM cells. **B)** HSCs had the highest total accessibility signal across all peaks of all hematopoietic cell types analyzed. Average cumulative signal across the master peak-list was determined by the -hist function of HOMER annotatePeaks.pl. **C-G)** Comparisons of the number of peaks gained and lost upon HSC differentiation into unipotent cells revealed that MkPs had the most similar accessibility profile to HSCs. **C)** Schematic of the pair-wise comparisons made. HSC peaks were compared with one unilineage cell type at a time and those comparisons are reported in D-G. **D)** MkPs had the lowest percentage of altered peaks from HSCs compared to the other 4 unilineage cell types. The percentage of all non-overlapping peaks (peaks both gained and lost) calculated as the ratio of unique peaks in each cell type when compared pairwise to HSCs divided by the total number of peaks called in that cell type are displayed here. The numbers in the bars represent the total number of peaks altered (gained+lost) for each cell type. EPs had the highest percentage of peaks altered (gained+lost), followed by T cells, GMs, then B cells. **E)** B cells had the highest percentage of peaks gained from HSCs, while MkPs had the lowest. Calculations as in **D)**, but only peaks gained are shown. **F)** EPs had the highest percentage of peaks lost from HSCs, while B cells had the least. Calculations as in **D)**, but only peaks lost are shown. **G)** B cells were the only lineage with more peaks gained (53%) than lost (47%) upon differentiation from HSCs. In this panel, the sum of peaks gained and lost in each cell type was set to 100% and then the ratio of peaks gained and lost was displayed. T cells had the second highest proportion of peaks gained (27%), followed by MkPs (19%), EPs (16%), then GMs (15%).

### Comparisons of peaks gained and lost as HSCs differentiate into unilineage cells revealed an overall gain of accessibility selectively for B cell differentiation

To assess the number of peaks that changed upon HSCs differentiation, we took the IDR optimal peak-list for each cell type and performed pair-wise comparisons between HSCs and the five mature/unipotent cell types (**Figure 3C**). We quantified the number of peaks gained and lost by the unipotent progenitors/mature cells compared to HSCs (**Figure 3C-G**). MkPs had the lowest number of peak changes (peaks gained plus lost; **Figure 3D**), and therefore have the greatest proportion of peaks in common with HSCs. This is primarily driven by the low percentage of peaks gained (**Figure 3E**), as opposed to peaks lost (**Figure 3F**) upon HSC differentiation into MkPs. In contrast, EPs had the highest percentage of total peaks changed (**Figure 3D**) due to the greatest percentage of peaks lost (**Figure 3F**). This could be driven by EPs starting to shut down transcription to become highly specialized and eject their nuclei, reflected by the overall low accessibility observed (**Figure 3A,B**). B cells had the highest percentage of peaks gained and the lowest percentage of peaks lost compared to the other cell types (**Figure 3E,F**) and was the only cell type where the percentage of peaks gained was higher than peaks lost (**Figure 3G**). This suggests that B cell fate requires chromatin remodeling to open up sites that drive B cell lineage fate.

### Exclusively shared peaks between HSCs and unipotent cell types are primarily non-promoter and are enriched for known cell-type specific transcription factors

We then turned our attention from peaks that were different between HSCs and their progeny to instead focus on elements with shared accessibility. We hypothesized that peaks that are exclusively shared between HSCs and one unipotent cell type contain elements that drive lineage commitment into that cell type. We filtered the peak lists of all 6 cell types against each other using the HOMER mergePeaks.pl tool and annotated the peak lists that each of unipotent lineage cell types exclusively shared with HSCs (**Figure 4A**). We quantified the percentage of peaks that each unipotent cell type shared with HSCs (**Figure 4B**). Consistent with the clustering profiles (**Figure 1C-D**), MkPs had the highest percentage of peaks that were shared exclusively with HSCs. This similarity appeared to be primarily manifested in non-promoter elements: we annotated the exclusively shared peaks and categorized them as promoter or non-promoter peaks (**Figure 4C**) and compared the distributions to the annotated peak lists for each cell type assayed (**Table 1**). All of the exclusively shared peak lists had significant enrichment (p-value <0.001) of non-promoter peaks compared to the normal distribution of peaks in our dataset. Thus, non-promoter elements are shared between HSCs and their progeny significantly more frequently than promoter elements, especially with MkPs.

**Figure 4:**
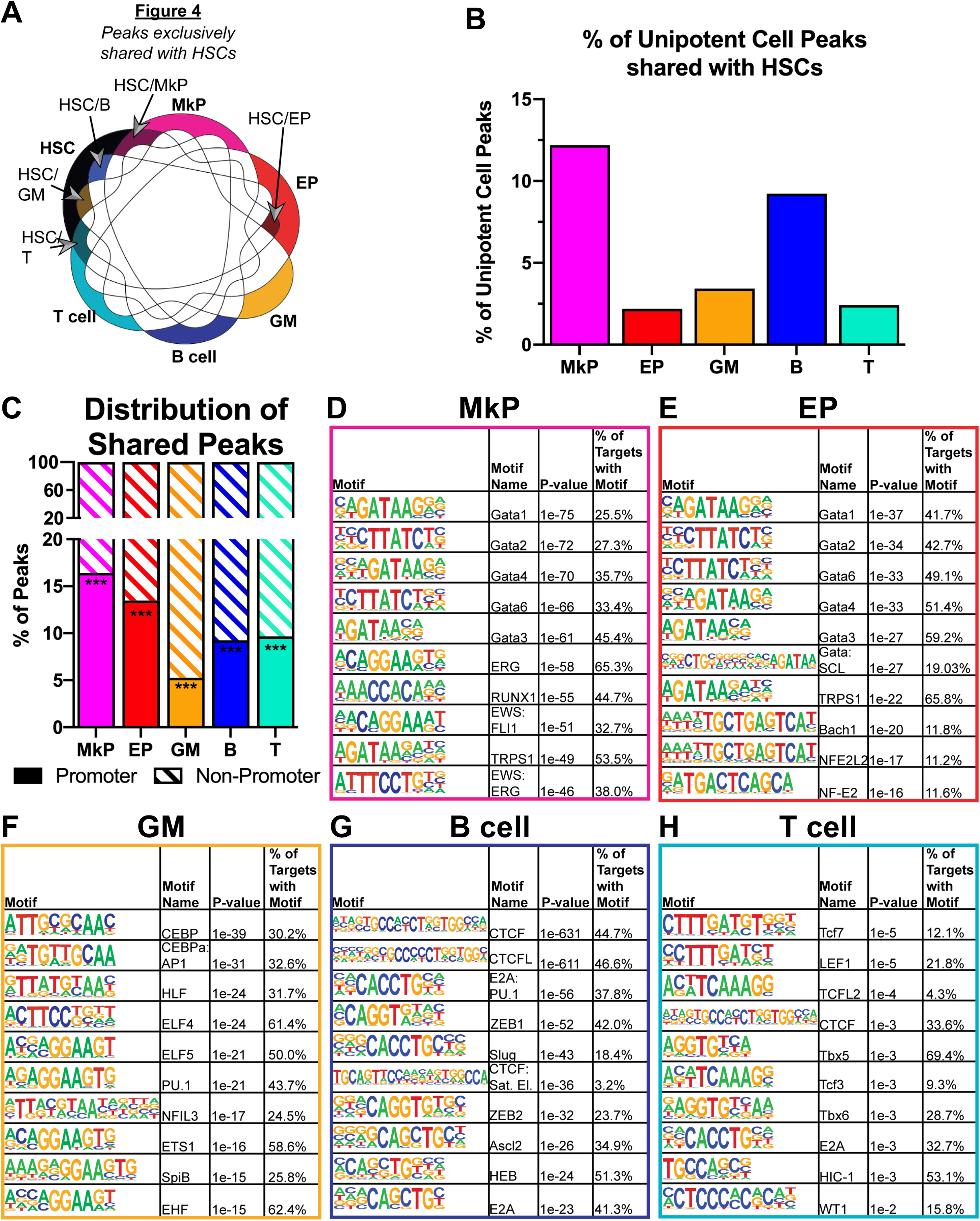
Peaks shared between HSCs and unipotent cell types are primarily non-promoter and are enriched for known cell type-specific transcription factors. **A)** Schematic for how the unipotent lineage peaks exclusively intersected with HSC peaks were generated. Peaks were compared using HOMER mergePeaks.pl tool using peaklists from the 6 cell types assayed. The resulting 5 overlapping peaklists contained shared peaks between HSCs and only the unipotent cell type of interest (but not present in any of the other four lineages). The five exclusive pairwise comparisons (e.g. HSC/MkP only, HSC/EP, etc.) were used for panels **B-H**. **B)** MkPs have the highest peak overlap with HSCs. The number of unipotent lineage peaks that were uniquely intersected with HSCs was divided by the total number of peaks for each mature cell type. MkPs had the highest percentage of HSC overlap (12.2%), followed by B cells (9.2%), GMs (3.4%), T cells (2.4%), then EPs (2.2%). **C)** Peaks exclusively shared between each unipotent cell type and HSCs were significantly enriched in the non-promoter regions of the genome. The shared peaklists described in **A)** were annotated using HOMER annotatePeaks.pl function and filtered as promoter (±500 bp from TSS), and non-promoter (< −500bp and > +500bp from TSS). The number of promoter and non-promoter peaks were divided by the total number of peaks for each cell type. For all cell types, less than 20% of peaks were promoter peaks, with MkPs with the highest (16.4%) and GMs with the lowest (5.3%) percentage. This is a significant (<0.001) difference compared to the normal distribution of promoter peaks (35-61%) for each cell type assayed. *** = p-value of <.001. **D-H)** Unipotent lineage peaks exclusively intersected with HSC peaks displayed enrichment of motifs for transcription factors with known roles in lineage differentiation. Motifs were found using HOMER findMotifsGenome.pl function, with a background file containing the combined peaklists of the other 4 cell types. The top 10 results, as ranked by p-value from the known_motifs.html output, are shown. **D)** In MkP/HSC peaks, Gata family peaks made up 5 of the top 10 hits, followed by ERG, Runx1, and fusions EWS:FL1 and EWS:ERG. **E)** EP/HSC-enriched motifs also contained Gata factors, as well as the combination Gata:SCL motif and the known beta-globin locus control binder NFE2 and its paralog NFE2L2. **F)** GM/HSCs had CEBPa and PU.1 motifs as top hits, along with ETS transcription factor binding sites. B cell/HSC-enriched motifs had CTCF with CTCFL (BORIS) as the top two hits. **G)** B cells/HSC peaks also had E2A motifs enriched, as well as Ascl2, Slug, and ZEB1/2. **H)** Tcf7 motif was the top hit for T cell/HSC-shared peaks, along with CTCF and Tbx5/6. Similar to the B-cell/HSC list, the T-cell/HSC list was also enriched for E2A motifs.

To determine what transcription factor binding sites were present within the exclusively shared peaks, we performed motif enrichment using the HOMER package and reported the top 10 results for each cell type, sorted by p-value (**Figure 4D-H**). The peaks that HSCs shared with MkPs (**Figure 4D**) or EPs (**Figure 4E**) were primarily enriched for Gata family transcription factors and their inhibitor TRPS1. Notably, HSC/MkP peaks also had enrichment of ERG and Runx1, which are known drivers of hematopoiesis (Growney et al., 2005; Kruse et al., 2009). For HSC/EPs, Gata1 was the most enriched motif, with the Gata:SCL combination motif and NF-E2 and NFE2L motifs also scoring in the top ten. These factors are all known to be important in red blood cell differentiation, and NF-E2 is known to regulate SCL and Gata2 (Siegwart et al., 2020). HSC/GM peaks had enrichment of known regulators of GM cell fate, such as CEBP, PU.1, and SpiB (**Figure 4F**). HSC/B cells primarily had CTCF and CTCFL motif enrichment (**Figure 4G**).

These motifs could be a reason for the overall high number of peaks observed in B cells (**Figure 3A,B**), as 44.7% and 46.6% of the shared peaks contained CTCF or CTCFL motifs, respectively. HSC/T cell peaks were enriched for Tcf and Tbx family factors that are known to play a role in T cell development (**Figure 4H**). Overall, all five HSC-shared peak lists had enrichment of transcription factors that are known to be important for normal differentiation for each lineage.

### Evidence of *cis* element priming of lineage-specific genes in HSCs

Previous work on understanding multipotency and developmental competence suggests a model where competence is conferred by transcriptional priming: being competent of transcription factor binding and gene expression, without active expression (Hu et al., 1997). One of the suggested regulators of transcriptional priming are *cis-*regulatory elements (CREs). This means that CREs that drive lineage fate for all lineages are accessible in HSCs in our permissive fate model and inaccessible in our *de novo* activation model. We hypothesized that CREs that are exclusively shared between HSCs and a unipotent lineage cell are potential drivers of that lineage cell. We utilized the GREAT tool (McLean et al., 2010) to annotate and predict the target genes for each exclusively shared CRE. Here we report examples of genes and a predicted CRE for each lineage that is primed in HSCs. In addition, we linked the motif enrichment with the GREAT analysis by annotating the CREs using the top 10 motifs enriched by p-value **(Figure 4D-H)** for each exclusive HSC/unipotent cell type. In MkPs, a predicted CRE for *Thrombin receptor like 2* (*F2rl2*) was found. This gene is expressed only in MkPs **(Figure 5A)**, while the CRE is only accessible in HSCs and MkPs **(Figure 5B)**. This CRE contained 9 out of the top 10 motifs, with the Runx1 motif being the only one missing **(Figure 5C)**. *Pyruvate kinase liver and red blood cell* (*Pklr)* was found to be expressed only in EPs **(Figure 5D)**, and a predicted CRE was accessible only in HSCs and EPs **(Figure 5E).** Motifs for Gata2, Gata3, Gata4, and TRPS1 were found within the CRE **(Figure 5F)**. In GMs, *Mitochondrial tumor suppressor 1* (*Mtus1*) was found to be primed in HSCs, with expression only in GMs **(Figure 5G)**, accessibility of a predicted CRE only in HSCs and GMs **(Figure 5H)**, and the presence of transcription factors known to play a role in GM development, such as CEBP and PU.1 **(Figure 5I)**. In B cells, *Interferon regulatory factor 8* (*Irf8*), is only expressed in B cells **(Figure 5J)**, the predicted CRE is only accessible in both B cells and HSCs **(Figure 5K)**, and contained 5 out of the top 10 motifs, ZEB1/2, Slug, Ascl2, HEB, and E2A **(Figure 5L)**. In T cells, the gene *Inducible T cell co-stimulator (Icos)* is only expressed in T cells **(Figure 5M)**, a predicted linked CRE is accessible in both T cells and HSCs **(Figure 5N)** and contains motifs for CTCF and WT1 **(Figure 5O)**. Taken together, these examples represent CRE priming in HSCs, along with the corresponding transcription factors that may act on each element to guide HSC fate.

**Figure 5:**
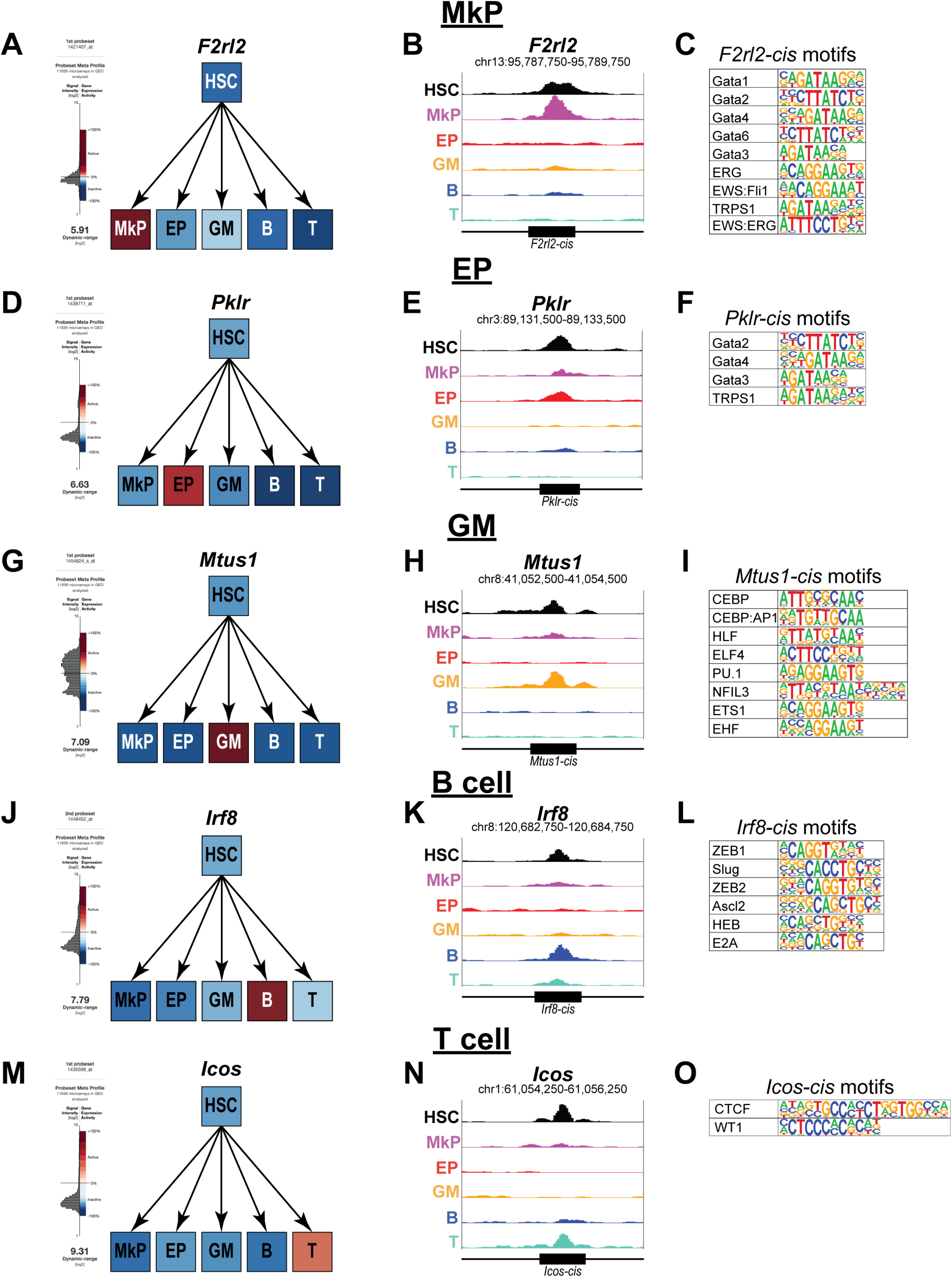
Examples of *cis* element priming of lineage-specific genes in HSCs. **A)** GEXC expression data reported expression of *Thrombin receptor like 2 (F2lr2)* selectively in MkPs. **B)** A *cis-*element predicted to be associated with *F2rl2* by GREAT was accessible in both MkPs and HSCs, but not in any other unipotent cell type. **C)** The *F2rl2* CRE contained the transcription factor binding motifs for 9 out of the top 10 enriched motifs in MkPs. The only motif not present is Runx1. **D)** GEXC expression data reported expression of *Pyruvate kinase liver and red blood cell (Pklr*) in EPs, and not any other cell type. **E)** A *cis-*element predicted to be associated with *Pklr* by GREAT was accessible in both EPs and HSCs, but not in any other unipotent cell type. **F)** The *Pklr* CRE contained the binding motifs for Gata2, Gata4, Gata3 and TRPS1. **G)** GEXC expression data reported selective expression of *Mitochondrial tumor suppressor 1* (*Mtus1*) in GMs and no expression in any other cell type. **H)** A *cis-*element predicted to be associated with *Mtus1* by GREAT was accessible in both GMs and HSCs. **I)** CEBP, CEBP:AP1, HLF, PU.1, NFL3, ETS1, and EHF binding motifs were present in the *Mtus1* CRE reported in (H). **J)** GEXC expression data reported *Interferon regulatory factor 8 (Irf8)* expression only in B cells, not in the other unipotent lineage cells or in HSCs. **K)** A *cis-*element predicted by GREAT to be associated with *Irf8* was accessible in both B cells and HSCs. **L)** ZEB1/2, Slug, Ascl2, HEB, and E2A binding motifs were found within the *Irf8* CRE displayed in (K). **M)** GEXC expression data reported *Inducible T cell co-stimulator (Icos)* expression only in T cells, but not in the other unipotent lineage cells or HSCs. **N)** A *cis-*element predicted by GREAT to be associated with *Icos* was accessible in both T cells and HSCs. **O)** CTCF and WT1 motifs were found within the *Icos* CRE displayed in (N).

## DISCUSSION

### MkPs and HSCs have the most similar accessibility profile

Here, we compared the genome-wide accessibility by ATAC-seq of the multipotent HSCs and unipotent lineage cell types (EPs, MkPs, GMs, B, and T cells). Through hierarchical clustering analysis, we observed erythromyeloid and lymphoid relationships that are consistent with the classical model of hematopoiesis **(Figure 1D)** (Boyer et al., 2011; Bryder et al., 2006; Laurenti and Göttgens, 2018; Seita and Weissman, 2010). By both PCA and hierarchical clustering, we observed that MkPs were the most similar to HSCs based on their accessibility profiles **(Figure 1)**. This relationship is reflected in a high level of overlap of peaks, as MkPs have the fewest peaks gained or lost from HSCs compared to the other cell types **(Figure 3)**and have the largest percentage of peaks exclusively shared with HSCs **(Figure 4B)**. These findings are in agreement with recent clonal studies of hematopoiesis that reported a megakaryocyte lineage bias of HSCs (Carrelha et al., 2018; Rodriguez-Fraticelli et al., 2018). According to hierarchal clustering, EPs had the second closest association to HSCs **(Figure 1D)** possibly supporting erythropoiesis as the default fate for hematopoiesis (Boyer et al., 2019) under conditions where chromatin remodeling silences megakaryocyte driver elements (Heuston et al., 2018). On the other end of the spectrum, the least similar cell types to HSCs were the lymphoid cell types **(Figure 1D)**. This greater difference was primarily due to a high proportion of peaks gained **(Figure 3E)**rather than lost **(Figure 3F)** upon differentiation from HSCs, leading to a greater ratio of peaks gained:lost for lymphoid cells than for erythromyeloid lineages **(Figure 3G)**.

### Evidence of multilineage priming in HSCs

The priming of genes for transcription likely initiates within CREs, which can then drive the activation of promoter targets. These enhancers can act as drivers of lineage fate (Wang et al., 2015) and their accessibility is a putative regulator of competence in stem cells. We made the assumption that peaks that are exclusively shared between HSCs and the unipotent lineage cells contain CREs that are specific for driving differentiation into that lineage. We observed that the majority of exclusively shared peaks were non-promoter peaks **(Figure 4B)**and were enriched for binding motifs of transcription factors known to be important for differentiation into each lineage **(Figure 4D-H)**. By using the GREAT tool, we made predictions for the target genes for the many CREs that were present in these exclusive lists. The examples shown in **Figure 5** provide evidence that multi-lineage priming exists in HSCs.

### Both permissive and *de novo* epigenetic mechanisms influence hematopoiesis

Analogous to other stem cell systems, multipotent HSCs with the competence to differentiate into diverse cell types reside at the top of the blood cell hierarchy. We tested two potential models of the mechanism of multipotency, the permissive fate and *de novo* activation **(Figure 1A)**. We found evidence for both. Supporting the permissive fate model are the observations that HSCs had the highest global accessibility **(Figure 3A/B)**, that peaks were lost in every unipotent cell type from HSCs **(Figure 3F)**, that every unipotent cell type shared some peaks exclusively with HSCs **(Figure 4B)**, and that evidence of multilineage priming of CREs were found in HSCs **(Figure 5)**. The *de novo* activation model was supported by the observation that new peaks were gained during differentiation into all five lineages (**Figure 3E**), and previous studies reporting progressive upregulation of lineage-specific genes as HSCs transition into progenitors (Forsberg et al., 2005; Terskikh et al., 2003). Thus, both mechanisms likely influence hematopoietic fate decisions. Interestingly, we found evidence that the balance between the two models varies between lineages. For example, B cells, and to a lesser extent T cells, had a higher proportion of peaks gained than lost compared to erythromyeloid lineages (**Figure 3G**). This may indicate that the megakaryocyte/erythroid lineage is in a more primed state in HSCs, whereas lymphopoiesis requires more extensive chromatin remodeling to both prime lymphoid CREs not accessible in HSCs and simultaneously shut down the megakaryocyte/erythrocyte trajectory. The cell output and kinetics from *in vivo* lineage tracing and reconstitution assays support these conclusions (Carrelha et al., 2018; Rodriguez-Fraticelli et al., 2018);(Boyer et al., 2011, 2012, 2019; Yamamoto et al., 2013). Our identification of specific, putative regulatory CREs will enable functional testing of these elements.

## EXPERIMENTAL PROCEDURES

### Mice and Cells

All experiments were performed using 8-to 12-week-old C57BL/6 wild-type mice in accordance with UCSC IACUC guidelines. Hematopoietic cells were isolated from BM by crushing murine femurs, tibias, hips, and sternums as previously described (Rajendiran et al., 2020). Stem and progenitor cell fractions were enriched using CD117-coupled magnetic beads (Miltenyi). Cells were stained with unconjugated lineage rat antibodies (CD3, CD4, CD5, CD8, B220, Gr1, Mac1, and Ter119) followed by goat-α-rat PE-Cy5 (Invitrogen). Stem and progenitor cells were isolated using fluorescently labeled or biotinylated antibodies for the following antigens: cKit (2B8, Biolegend), Sca1 (D7, Biolegend), Slamf1(CD150) (TC15-12F12.2, Biolegend), CD41(MWReg30, Biolegend), and CD71(RI7217, Biolegend). Cells were sorted using a FACS Aria II (BD Bioscience). HSCs were defined as cKit^+^ Lin^−^ Sca1^+^ Flk2^−^ and Slamf1^+^; MkPs as cKit^+^Lin^−^Sca1^−^Slamf1^−^CD41^+^. Unipotent lineage cells were isolated by the following markers and as described previously (Cool et al., 2020; Leung et al., 2019) : EPs, Lin(CD3, CD4, CD5, CD8, B220, Gr1, and Mac1)^−^ CD71^+^Ter119^+/−^; GMs, Lin(CD3, CD4, CD5, CD8, B220, and Ter119)^−^ Gr1^+^Mac1^+^ (“GM” cells were positive for both Gr1 and Mac1); T cells, Lin(CD5, B220, Gr1, Mac1, and Ter119)^−^ CD25^−^CD3^+^CD4^+/−^CD8^+/−^; B cells, Lin(CD3, CD4, CD8, Gr1, Mac1, and Ter119)^−^ CD43^−^B220^+^.

### ATAC-seq

ATAC-seq was performed as previously described (Buenrostro et al., 2013). Briefly, cells were collected after sorting into microcentrifuge tubes containing staining media (1xDPBS,1mM EDTA with 5% serum). They were centrifuged at 500xg for 5 minutes at 4°C to pellet the cells. The supernatant was aspirated, and the cells were washed with ice-cold 1xDPBS. Cells were centrifuged and the supernatant was discarded. Cells were immediately resuspended in ice-cold lysis buffer (10 mM Tris-HCl, pH 7.4, 10 mM NaCl, 3 mM MgCl2 and 0.1% IGEPAL CA-630) and centrifuged at 500xg for 10 minutes. The supernatant was aspirated, and pellets were resuspended in transposase reaction mix (25μL 2xTD Buffer, 2.5μL transposase (Illumina), and 22.5μL nuclease free water). The transposition reaction was carried out at 37°C for 30 minutes at 600rpm in a shaking thermomixer (Eppendorf). Immediately after completion of the transposition reaction, the samples were purified using the MinElute Reaction Clean up kit (Qiagen) and eluted into 10 μL of EB. Samples were stored at −20°C until PCR amplification step. PCR amplification was performed as previously described (Buenrostro et al., 2013) using custom Nextera primers. After initial amplification, a portion of the samples were run on qPCR (ViiA7 Applied Biosystems) to determine the additional number of cycles needed for each library. The libraries were purified using the MinElute Reaction Clean up kit (Qiagen), eluted into 20 μL EB and then size selected using AmpureXP(Beckman-Coulter) beads at a ratio of 1.8:1 beads/sample, and eluted into 40μL of nuclease-free water. Library size distribution was determined by Bioanalyzer (Agilent) capillary electrophoresis and library concentration was determined by Qubit 3 (Life Technologies). Quality of libraries were checked by shallow sequencing (1 million raw reads) on a Miseq (Illumina) at 75 x 75 paired-end sequencing. Those libraries that appeared to have size distributions similar to previous reports were pooled together and deep sequenced on a HiSeq2500 (Illumina) at 100 x 100 reads at the Vincent J. Coates Genomics Sequencing Laboratory at UC Berkeley.

### Data processing

Demultiplexed sequencing data was processed using the ENCODE ATAC-seq pipeline version 1.1.6 and 1.4.2 (https://github.com/ENCODE-DCC/atac-seq-pipeline) using the mm10 assembly and the default parameters. In version 1.4.2 changed: atac.multimapping=0, atac.smooth_win=150, atac.enable_idr=true, atac.idr_thresh=0.1 to be consistent with the mapping/peak calling performed with previous versions.

Peak filtering, hierarchical clustering, and tSNE plot production was performed using the chromVAR package (https://github.com/GreenleafLab/chromVAR). First, the optimal peak-list from the IDR output for each cell type was concatenated and sorted, then used as the peak input for chromVAR. The blacklist filtered bam files for reach replicate was used as input along with the sorted peak file. The fragment counts in each peak for each replicate and GC bias was calculated, and then the peaks were filtered using filterPeaks function with the default parameters and nonoverlapping=TRUE. The master peak-list was extracted at this point, which contained 84,243 peaks, and used throughout the study. The deviations were calculated using every peak, and the tSNE and correlation functions were also performed using the deviations output and the default parameters.

Annotation of peaks, generation of histogram plot, merging of peaks, and motif enrichment was performed by HOMER (http://homer.ucsd.edu/homer/). Peaks were annotated using the annotatePeaks.pl function with the mm10 assembly and default parameters. Histogram was created by first shifting the bam files using DeepTools alignmentSieve.py with the flag –ATACshift. Next, tag directories were made using the Tn5 shifted bam files using HOMER makeTagDirectory. The histogram was made using the annotatePeaks.pl function with the default settings and the flags: -size −500,500 and -hist 5. Peak lists were compared using the mergePeaks.pl function with default settings and the flags -d given, -venn, and for the unique peak lists -prefix. Motif enrichment was performed using the findMotifsGenome.pl package with default parameters using the flag -size given and custom background peaks, which consisted of the combination of all the peaklists for the cell types not being analyzed. Instances of motifs in non-promoter peaks were found by using the annotatePeaks.pl function with the -m flag, using custom made motif files for each cell type containing the top 10 enriched motifs found.

The GREAT tool (http://great.stanford.edu/public/html/) was used to annotate non-promoter peaks to target genes. The peak lists were reduced to BED4 files from the HOMER annotations output and used as input. The whole mm10 genome was used as the background regions, and the association rule settings were set as Basal plus extension, proximal window 2kb upstream, 1kb downstream, plus distal up to 1Mb and included curated regulatory domains. All genome track visualizations were made using the UCSC genome browser. Graphs were made in either Microsoft Excel or GraphPad Prism 8. Annotations to figures was performed using Adobe Illustrator CC and Adobe Photoshop CC.

## Authors’ Contributions

E.W.M., J.K., and E.C.F. designed the experiments. E.W.M., J.K, and R.S. isolated and sorted the primary cell types and performed ATAC-seq. E.W.M and R.E.R. conducted data processing and analysis. E.W.M. and E.C.F. wrote the paper. All authors reviewed the manuscript.

## Acknowledgements

We thank Bari Nazario and the IBSC flow cytometry core for assistance and support; Sol Katzman for bioinformatic assistance; and Forsberg lab members for comments on the manuscript. The authors declare that they have no competing interests.

## REFERENCES

An, X., Schulz, V.P., Li, J., Wu, K., Liu, J., Xue, F., Hu, J., Mohandas, N., and Gallagher, P.G. (2014). Global transcriptome analyses of human and murine terminal erythroid differentiation. Blood 123, 3466–3477.

Bernstein, B.E., Mikkelsen, T.S., Xie, X., Kamal, M., Huebert, D.J., Cuff, J., Fry, B., Meissner, A., Wernig, M., Plath, K., et al. (2006). A Bivalent Chromatin Structure Marks Key Developmental Genes in Embryonic Stem Cells. Cell 125, 315–326.

Boyer, S.W., Schroeder, A. V., Smith-Berdan, S., and Forsberg, E.C. (2011). All Hematopoietic Cells Develop from Hematopoietic Stem Cells through Flk2/Flt3-Positive Progenitor Cells. Cell Stem Cell 9, 64–73.

Boyer, S.W., Beaudin, A.E., and Forsberg, E.C. (2012). Mapping differentiation pathways from hematopoietic stem cells using Flk2/Flt3 lineage tracing. Cell Cycle 11, 3180–3188.

Boyer, S.W., Rajendiran, S., Beaudin, A.E., Smith-berdan, S., Muthuswamy, P.K., Perez-Cunningham, J., Martin, E.W., Cheung, C., Tsang, H., Landon, M., et al. (2019). Clonal and Quantitative In Vivo Assessment of Hematopoietic Stem Cell Differentiation Reveals Strong Erythroid Potential of Multipotent Cells. Stem Cell Reports 12, 801–815.

Bryder, D., Rossi, D.J., and Weissman, I.L. (2006). Hematopoietic Stem Cells: The Paradigmatic Tissue-Specific Stem Cell. Am. J. Pathol. 169, 338–346.

Buenrostro, J.D., Giresi, P.G., Zaba, L.C., Chang, H.Y., and Greenleaf, W.J. (2013). Transposition of native chromatin for fast and sensitive epigenomic profiling of open chromatin, DNA-binding proteins and nucleosome position. Nat Meth 10, 1213–1218.

Buenrostro, J.D., Corces, M.R., Lareau, C.A., Wu, B., Schep, A.N., Aryee, M.J., Majeti, R., Chang, H.Y., and Greenleaf, W.J. (2018). Integrated Single-Cell Analysis Maps the Continuous Regulatory Landscape of Human Hematopoietic Differentiation. Cell 0, 1–14.

Bulut-Karslioglu, A., Macrae, T.A., Oses-Prieto, J.A., Covarrubias, S., Percharde, M., Ku, G., Diaz, A., McManus, M.T., Burlingame, A.L., and Ramalho-Santos, M. (2018). The Transcriptionally Permissive Chromatin State of Embryonic Stem Cells Is Acutely Tuned to Translational Output. Cell Stem Cell.

Cabal-Hierro, L., van Galen, P., Prado, M.A., Higby, K.J., Togami, K., Mowery, C.T., Paulo, J.A., Xie, Y., Cejas, P., Furusawa, T., et al. (2020). Chromatin accessibility promotes hematopoietic and leukemia stem cell activity. Nat. Commun. 11, 1406.

Cabezas-Wallscheid, N., Klimmeck, D., Hansson, J., Lipka, D.B., Reyes, A., Wang, Q., Weichenhan, D., Lier, A., Von Paleske, L., Renders, S., et al. (2014). Identification of regulatory networks in HSCs and their immediate progeny via integrated proteome, transcriptome, and DNA methylome analysis. Cell Stem Cell 15, 507–522.

Carrelha, J., Meng, Y., Kettyle, L.M., Luis, T.C., Norfo, R., Alcolea, V., Boukarabila, H., Grasso, F., Gambardella, A., Grover, A., et al. (2018). Hierarchically related lineage-restricted fates of multipotent haematopoietic stem cells. Nature 554, 106–111.

Cool, T., Worthington, A., Poscablo, D., Hussaini, A., and Forsberg, E.C. (2020). Interleukin 7 receptor is required for myeloid cell homeostasis and reconstitution by hematopoietic stem cells. Exp. Hematol. Oct;90:39–45.e3. doi: 10.1016/j.exphem.2020.09.001

Forsberg, E.C., Downs, K.M., Christensen, H.M., Im, H., Nuzzi, P.A., and Bresnick, E.H. (2000). Developmentally dynamic histone acetylation pattern of a tissue-specific chromatin domain. Proc. Natl. Acad. Sci. 97, 14494 LP–14499.

Forsberg, E.C., Prohaska, S.S., Katzman, S., Heffner, G.C., Stuart, J.M., and Weissman, I.L. (2005). Differential expression of novel potential regulators in hematopoietic stem cells. PLoS Genet. 1.

Gaspar-Maia, A., Alajem, A., Polesso, F., Sridharan, R., Mason, M.J., Heidersbach, A., Ramalho-Santos, J., McManus, M.T., Plath, K., Meshorer, E., et al. (2009). Chd1 regulates open chromatin and pluripotency of embryonic stem cells. Nature 460, 863–868.

Gaspar-Maia, A., Alajem, A., Meshorer, E., and Ramalho-Santos, M. (2011). Open chromatin in pluripotency and reprogramming. Nat. Rev. Mol. Cell Biol. 12, 36–47.

Growney, J.D., Shigematsu, H., Li, Z., Lee, B.H., Adelsperger, J., Rowan, R., Curley, D.P., Kutok, J.L., Akashi, K., Williams, I.R., et al. (2005). Loss of Runx1 perturbs adult hematopoiesis and is associated with a myeloproliferative phenotype. Blood 106, 494–504.

Heinz, S., Benner, C., Spann, N., Bertolino, E., Lin, Y.C., Laslo, P., Cheng, J.X., Murre, C., Singh, H., and Glass, C.K. (2010). Simple Combinations of Lineage-Determining Transcription Factors Prime cis-Regulatory Elements Required for Macrophage and B Cell Identities. Mol. Cell 38, 576–589.

Hestdal, K., Ruscetti, F.W., Ihle, J.N., Jacobsen, S.E., Dubois, C.M., Kopp, W.C., Longo, D.L., and Keller, J.R. (1991). Characterization and regulation of RB6-8C5 antigen expression on murine bone marrow cells. J. Immunol. 147, 22 LP–28.

Heuston, E.F., Keller, C.A., Lichtenberg, J., Giardine, B., Anderson, S.M., Hardison, R.C., and Bodine, D.M. (2018). Establishment of regulatory elements during erythro-megakaryopoiesis identifies hematopoietic lineage-commitment points. Epigenetics and Chromatin 11, 1–18.

Hu, M., Krause, D., Greaves, M., Sharkis, S., Dexter, M., Heyworth, C., and Enver, T. (1997). Multilineage gene expression precedes commitment in the hemopoietic system. Genes Dev. 11, 774–785.

Kruse, E.A., Loughran, S.J., Baldwin, T.M., Josefsson, E.C., Ellis, S., Watson, D.K., Nurden, P., Metcalf, D., Hilton, D.J., Alexander, W.S., et al. (2009). Dual requirement for the ETS transcription factors Fli-1 and Erg in hematopoietic stem cells and the megakaryocyte lineage. Proc. Natl. Acad. Sci. 106, 13814 LP–13819.

Laurenti, E., and Göttgens, B. (2018). From haematopoietic stem cells to complex differentiation landscapes. Nature 553, 418–426.

Leung, G.A., Cool, T., Valencia, C.H., Worthington, A., Beaudin, A.E., and Camilla Forsberg, E. (2019). The lymphoid-associated interleukin 7 receptor (IL7R) regulates tissue-resident macrophage development. Dev. 146, dev176180.

Li, Q., Brown, J.B., Huang, H., and Bickel, P.J. (2011). Measuring reproducibility of high-throughput experiments. Ann. Appl. Stat. 5, 1752–1779.

Månsson, R., Hultquist, A., Luc, S., Yang, L., Anderson, K., Kharazi, S., Al-Hashmi, S., Liuba, K., Thorén, L., Adolfsson, J., et al. (2007). Molecular Evidence for Hierarchical Transcriptional Lineage Priming in Fetal and Adult Stem Cells and Multipotent Progenitors. Immunity 26, 407–419.

McLean, C.Y., Bristor, D., Hiller, M., Clarke, S.L., Schaar, B.T., Lowe, C.B., Wenger, A.M., and Bejerano, G. (2010). GREAT improves functional interpretation of cis-regulatory regions. Nat. Biotechnol. 28, 495–501.

Rajendiran, S., Smith-Berdan, S., Kunz, L., Risolino, M., Selleri, L., Schroeder, T., and Forsberg, E.C. (2020). Ubiquitous overexpression of CXCL12 confers radiation protection and enhances mobilization of hematopoietic stem and progenitor cells. Stem Cells 38, 1159–1174.

Rodriguez-Fraticelli, A.E., Wolock, S.L., Weinreb, C.S., Panero, R., Patel, S.H., Jankovic, M., Sun, J., Calogero, R.A., Klein, A.M., and Camargo, F.D. (2018). Clonal analysis of lineage fate in native haematopoiesis. Nature 553, 212–216.

Schep, A.N., Wu, B., Buenrostro, J.D., and Greenleaf, W.J. (2017). ChromVAR: Inferring transcription-factor-associated accessibility from single-cell epigenomic data. Nat. Methods 14, 975–978.

Seita, J., and Weissman, I.L. (2010). Hematopoietic stem cell: self-renewal versus differentiation. WIREs Syst. Biol. Med. 2, 640–653.

Seita, J., Sahoo, D., Rossi, D.J., Bhattacharya, D., Serwold, T., Inlay, M.A., Ehrlich, L.I.R., Fathman, J.W., Dill, D.L., and Weissman, I.L. (2012). Gene expression commons: An open platform for absolute gene expression profiling. PLoS One 7, 1–11.

Shivdasani, R.A., Fujiwara, Y., McDevitt, M.A., and Orkin, S.H. (1997). A lineage-selective knockout establishes the critical role of transcription factor GATA-1 in megakaryocyte growth and platelet development. EMBO J. 16, 3965–3973.

Siegwart, L.C., Schwemmers, S., Wehrle, J., Koellerer, C., Seeger, T., Gründer, A., and Pahl, H.L. (2020). The transcription factor NFE2 enhances expression of the hematopoietic master regulators SCL/TAL1 and GATA2. Exp. Hematol. 87, 42–47.e1.

Smith-Berdan, S., Bercasio, A., Rajendiran, S., and Forsberg, E.C. (2019). Viagra Enables Efficient, Single-Day Hematopoietic Stem Cell Mobilization. Stem Cell Reports 13, 787–792.

Terskikh, A. V, Miyamoto, T., Chang, C., Diatchenko, L., and Weissman, I.L. (2003). Gene expression analysis of purified hematopoietic stem cells and committed progenitors. Blood 102, 94–101.

Ugarte, F., Sousae, R., Cinquin, B., Martin, E.W., Krietsch, J., Sanchez, G., Inman, M., Tsang, H., Warr, M., Passegué, E., et al. (2015). Progressive Chromatin Condensation and H3K9 Methylation Regulate the Differentiation of Embryonic and Hematopoietic Stem Cells. Stem Cell Reports 5, 728–740.

Waddington, C.H. (1940). Organisers and genes (University Press; Cambridge).

Wang, A., Yue, F., Li, Y., Xie, R., Harper, T., Patel, N.A., Muth, K., Palmer, J., Qiu, Y., Wang, J., et al. (2015). Epigenetic priming of enhancers predicts developmental competence of hESC-derived endodermal lineage intermediates. Cell Stem Cell 16, 386–399.

Xu, J., Watts, J.A., Pope, S.D., Gadue, P., Kamps, M., Plath, K., Zaret, K.S., and Smale, S.T. (2009). Transcriptional competence and the active marking of tissue-specific enhancers by defined transcription factors in embryonic and induced pluripotent stem cells. Genes Dev. 23, 2824–2838.

Yamamoto, R., Morita, Y., Ooehara, J., Hamanaka, S., Onodera, M., Rudolph, K.L., Ema, H., and Nakauchi, H. (2013). Clonal Analysis Unveils Self-Renewing Lineage-Restricted Progenitors Generated Directly from Hematopoietic Stem Cells. Cell 154, 1112–1126.

